# The ERK MAPK pathway modulates Gq-dependent locomotion in *>Caenorhabditis elegans*

**DOI:** 10.1101/231746

**Authors:** Brantley Coleman, Irini Topalidou, Michael Ailion

## Abstract

The heterotrimeric G protein Gq regulates neuronal activity through distinct downstream effector pathways. In addition to the canonical Gq effector phospholipase Cβ, the small GTPase Rho was recently identified as a conserved effector of Gq. To identify additional molecules important for Gq signaling in neurons, we performed a forward genetic screen in the nematode *Caenorhabditis elegans* for suppressors of the hyperactivity and exaggerated waveform of an activated Gq mutant. We isolated two mutations affecting the MAP kinase scaffold protein KSR-1 and found that KSR-1 modulates locomotion downstream of or in parallel to the Gq-Rho pathway. Through epistasis experiments, we found that the core ERK MAPK cascade is required for Gq-Rho regulation of locomotion, but that the canonical ERK activator LET-60/Ras may not be required. Through neuron-specific rescue experiments, we found that the ERK pathway functions in head acetylcholine neurons to control Gq-dependent locomotion. Additionally, expression of activated LIN-45/Raf in head acetylcholine neurons is sufficient to cause an exaggerated waveform phenotype and hypersensitivity to the acetylcholinesterase inhibitor aldicarb, similar to an activated Gq mutant. Taken together, our results suggest that the ERK MAPK pathway modulates the output of Gq-Rho signaling to control locomotion behavior in *C. elegans*.

## Introduction

The heterotrimeric G protein Gq is a conserved regulator of neurotransmission in metazoans. Gq is highly expressed in neurons in mammals and in the nematode *C. elegans* (Wilkie *et al.* 1991; Lackner *et al.* 1999). In its canonical signaling pathway, Gq activates phospholipase Cβ (PLCβ) to cleave the membrane lipid phosphatidylinositol 4,5-bisphosphate (PIP2) into inositol trisphosphate (IP3) and diacylglycerol (DAG) (Rhee 2001). An increased DAG concentration at the synapse helps trigger synaptic vesicle release (Miller *et al.* 1999; Lackner *et al.* 1999).

In addition to activating PLCβ, Gq directly binds and activates the Rho guanine nucleotide exchange factor (GEF) Trio, which in turn activates the small GTPase Rho (Lutz *et al.* 2005, 2007; Williams *et al.* 2007). In mature *C. elegans* neurons, the Rho ortholog RHO-1 regulates synaptic activity through multiple G protein-dependent mechanisms. First, RHO-1 acts downstream of the G_12_-class G protein GPA-12 by binding to and inhibiting the diacylglycerol kinase DGK-1. Inhibition of DGK-1 allows DAG to accumulate at the synapse, thereby increasing synaptic vesicle release (McMullan *et al.* 2006; Hiley *et al.* 2006). Second, Gq-Rho signaling promotes neurotransmitter release by recruiting the sphingosine kinase SPHK-1 to presynaptic terminals (Chan *et al.* 2012). Third, Gq-Rho signaling positively regulates the NCA-1/NALCN cation channel to regulate locomotion (Topalidou *et al.* 2017a). Here we identify the extracellular signal-related kinase mitogen-activated protein kinase (ERK MAPK) pathway as a positive regulator of neuronal activity acting downstream of or in parallel to Gq and Rho in acetylcholine neurons.

ERK MAPK signaling acts extensively in animal development, cellular proliferation, and cancer signaling (Yoon and Seger 2006; Karnoub and Weinberg 2008; Sun *et al.* 2015). ERKs are highly expressed in mammalian neurons (Boulton *et al.* 1991; Ortiz *et al.* 1995) and act in both the nucleus and at the synapse to regulate synaptic activity and plasticity (Thomas and Huganir 2004; Sweatt 2004; Mao and Wang 2016b). In *C. elegans*, the ERK pathway is required for multiple developmental events including specification of the vulva (Sternberg 2005; Sundaram 2013), and also acts in several types of neurons to control behavior. ERK signaling is activated in response to odorants in the AWC sensory neuron to regulate chemotaxis to volatile odorants and in AIY interneurons to mediate odor adaptation (Hirotsu *et al.* 2000; Hirotsu and Iino 2005; Chen *et al.* 2011; Uozumi *et al.* 2012). ERK is also activated in the ASER sensory neuron to regulate chemotaxis to salt (Tomioka *et al.* 2006; Tomida *et al.* 2012). ERK signaling regulates foraging behavior by acting in the IL1, OLQ, and RMD neurons (Hamakawa *et al.* 2015). Finally, the ERK pathway has been shown to act in interneurons to regulate the nose touch response, a mechanosensory behavior (Hyde *et al.* 2011).

In the canonical ERK MAPK pathway, extracellular ligand binding activates transmembrane receptor tyrosine kinases (RTKs), and adaptor proteins recruit a GEF to activate the small GTPase Ras (LET-60 in *C. elegans*). Upon Ras activation, LIN-45/Raf translocates to the plasma membrane where it interacts with Ras and the scaffold protein KSR-1. KSR-1 facilitates the activation of LIN-45/Raf and the subsequent phosphorylation of the MAPK cascade consisting of LIN-45/Raf, MEK-2/MEK, and MPK-1/ERK (Sundaram 2013). In this study, we found that the ERK MAPK pathway consisting of KSR-1, LIN-45/Raf, MEK-2/MEK and MPK-1/ERK modulates Gq-Rho signaling in acetylcholine neurons, but that surprisingly LET-60/Ras may not be required.

## Materials and Methods

### C. elegans strains

All strains were cultured using standard methods and were maintained at 20°C. Table S1 contains all the strains used in this study.

### Isolation and mapping of the ksr-1(ox314) and ksr-1(yak10) mutations

The *ox314* and *yak10* mutants were isolated from an ENU mutagenesis suppressor screen of the activated Gq mutant *egl-30(tg26)* (Ailion *et al.* 2014). We mapped the *ox314* mutation by its activated Gq suppression phenotype using single nucleotide polymorphisms (SNPs) in the Hawaiian strain CB4856 as described (Davis *et al.* 2005).The *ox314* mutation was mapped to an approximately 709 kb region in the middle of the X chromosome between SNPs on cosmids F45E1 and F53A9 (SNPs F45E1[1] and pkP6158). This region included 159 predicted protein-coding genes. A complementation test of *ox314* and *yak10* in the *egl-30(tg26)* background showed these to be alleles of the same gene. Whole genome sequencing (see below) identified these as mutations in *ksr-1*, and we confirmed this by performing a complementation test with the deletion allele *ksr-1(ok786)*.

### Whole genome sequencing

Strains EG4198 *egl-30(tg26); ox314* and XZ1340 *egl-30(tg26); yak10* were sequenced to identify candidate mutations. DNA was purified according to the Hobert Lab protocol (http://hobertlab.org/whole-genome-sequencing/). Ion Torrent sequencing was performed at the University of Utah DNA Sequencing Core Facility. Each data set contained roughly 18,400,000 reads of a mean read length of 160 bases, resulting in about 30X average coverage of the *C. elegans* genome. The sequencing data were processed on the Galaxy server at usegalaxy.org (Afgan *et al.* 2016). SNPs and indels were identified and annotated using the Unified Genotyper and SnpEff tools (DePristo *et al.* 2011; Cingolani *et al.* 2012). After filtering for mutations in open reading frames, we found each strain to have unique stop mutations in *ksr-1*, in the middle of the interval where we mapped *ox314*. *ox314* is a G to A transition that causes a stop codon at amino acid K463, and *yak10* is an A to T transversion that causes a stop codon at W254.

### Locomotion assays

Track waveform and radial locomotion assays were performed on 10 cm nematode growth medium (NGM) plates seeded with 400 µl of OP50 *E. coli* culture and spread with sterile glass beads. Bacterial lawns were grown at 37°C for 16 hrs and the plates were stored at 4°C until needed. For track waveform measurements, five first day adult animals were placed on a plate and allowed to roam for 2-5 min. We then recorded each animal’s tracks following forward locomotion. Track pictures were taken at 40X on a Nikon SMZ18 microscope with the DS-L3 camera control system. Pictures of worm tracks were processed using ImageJ. Period and 2X amplitude were measured freehand using the line tool. For each worm, we calculated the average period/amplitude ratio of five individual track bends (Figure 1C). For assays with the temperature sensitive allele *sos-1(cs41)*, all strains were grown at 20°C and shifted to the non-permissive temperature of 25°C for 24 hours before being assayed. For radial locomotion assays, ten to fifteen first day adult animals were picked to the center of a plate and were then allowed to move freely for 40 minutes. The positions of worms were marked and the distances of the worms from the starting point were measured. For all waveform and radial locomotion assays, the experimenter was blind to the genotypes of the strains assayed.

**Figure 1.**
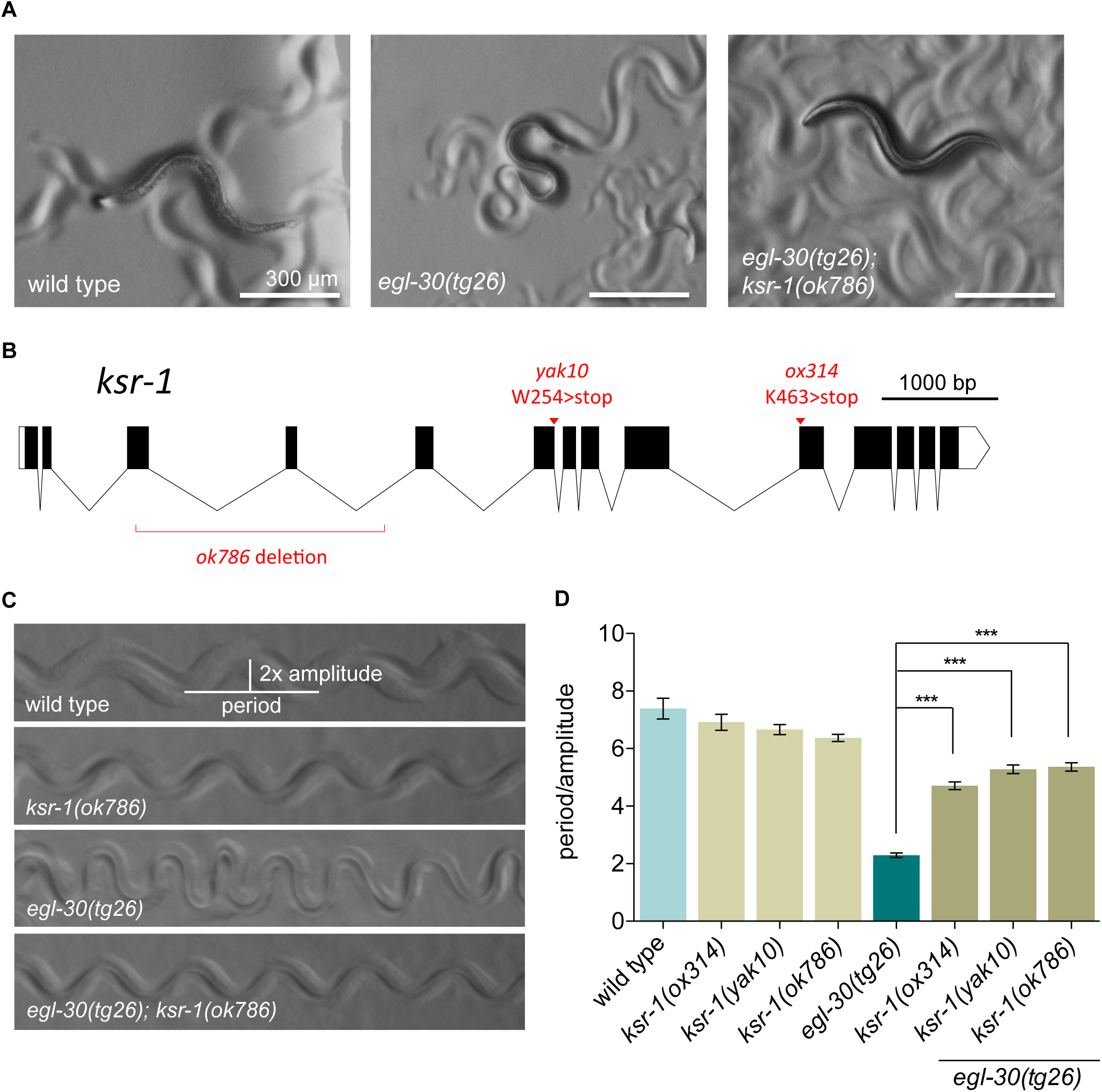
*ksr-1* mutations suppress activated Gq. (A) *ksr-1* mutation suppresses the coiled posture of activated Gq. The activated Gq mutant *egl-30(tg26)* has small size and deep body bends. *ksr-1(ok786)* suppresses the exaggerated body bends and small size of *egl-30(tg26)* worms. (B) Gene structure of *ksr-1* locus. Locations of the *egl-30(tg26)* suppressor alleles *ox314* and *yak10* are indicated, as well as the position of the *ok786* deletion. The gene structure was drawn using Exon-Intron Graphic Maker (www.wormweb.org/exonintron) (C) *ksr-1* mutation suppresses the exaggerated waveform of *egl-30(tg26)* mutants. Straightened images of tracks left in bacterial lawns show similar waveform for wild type and *ksr-1(ok786)* worms. *egl-30(tg26)* mutants have an exaggerated waveform, creating tracks with a large amplitude relative to the period. *ksr-1*(*ok786)* suppresses the exaggerated waveform of the *egl-30(tg26)* mutant. (D) *ksr-1* mutations suppress the *egl-30(tg26)* exaggerated waveform. *ksr-1* nonsense alleles *ox314* and *yak10* and the deletion allele *ok786* strongly but incompletely suppress the *egl-30(tg26)* exaggerated waveform. *N ≥* 12, *** *P* < 0.001, one-way ANOVA with Bonferroni’s *post hoc* test. All *egl-30(tg26); ksr-1* double mutants were significantly different than *ksr-1* single mutants (P<0.05), indicating that suppression is incomplete.

### Microscopy

Photographs of moving worms were taken at 60X on a Nikon SMZ18 microscope with the DS-L3 camera control system. The worms were age-matched as first day adults grown at 20°C.

### Constructs and transgenes

Plasmids were constructed using the three-slot multisite Gateway cloning system (Invitrogen). Plasmids and primers used are found in Tables S2 and S3. The *ksr-1*, *lin-45*, *let-60*, and *egl-30* cDNAs were amplified by PCR from a worm cDNA library and cloned into [1-2] Gateway entry vectors. Activating Raf mutations T626E/T629D were introduced into the *lin-45* cDNA vector by two sequential site-directed mutagenesis reactions (Q5 kit, NEB) with primers oBC094/095 and oBC096/097, respectively, and then confirmed by sequencing. The dominant Ras mutation S17N was introduced into the *let-60* cDNA by site-directed mutagenesis with primers oBC109/110. The *tg26* activating Gq mutation R243Q was introduced into the *egl-30* cDNA by site-directed mutagenesis with primers oET254/255. cDNAs were cloned into expression constructs under different neuronal promoters using the multisite Gateway system. Specifics of promoter fragments used are given in Table S2. For expression in different subclasses of acetylcholine neurons, we used the *Punc-17H* promoter that is expressed in head acetylcholine neurons (Hammarlund *et al.* 2007; Topalidou *et al.* 2017a) and the *Punc-17β* promoter that is expressed in ventral cord acetylcholine motor neurons (Charlie *et al.* 2006; Topalidou *et al.* 2017a). Proper expression of *ksr-1*, *lin-45* and *let-60* was confirmed by including an operon GFP::H2B in the [2-3] slot of the expression constructs. The operon GFP template *tbb-2 3’utr::gpd-2 operon::GFP::H2B:cye-1 3’utr* (Frøkjær-Jensen *et al.* 2012) results in untagged proteins whose expression can be monitored by GFP expression.

### Injections and chromosomal integrations

Worms carrying the activated *lin-45* transgenes *Punc-17::lin-45\** and *Punc-17H::lin-45\** as extrachromosomal arrays were generated by injecting pBC37 or pBC44 at 20 ng/µL or 10 ng/µL respectively along with co-injection markers pCFJ104 (*Pmyo-3::mCherry*) at 5 ng/µL, pCFJ90 (*Pmyo-2::mCherry*) at 2.5 ng/µL, and the carrier DNA Litmus 38i to a final concentration of 100 ng/µL DNA (Mello *et al.* 1991). Worms carrying the dominant negative *let-60* transgene *Prab-3::let-60(S17N)* were generated by injecting pBC50 at 20 ng/ul with the same co-injection markers. Worms carrying the activated Gq *egl-30(tg26)* transgene *Punc-17H::egl-30(tg26)* were generated by injecting pET102 at 10 ng/ul with the co-injection marker pCFJ90 (*Pmyo-2::mCherry*) at 1 ng/µL. MosSCI lines were generated as described (Frøkjær-Jensen *et al.* 2012) using an injection mix containing 10-15 ng/µL targeting vector, 50 ng/µL pCFJ601 (*Peft-3::Mos1* transposase), negative selection markers pGH8 (*Prab-3::mCherry*) at 10 ng/µL, pCFJ104 (*Pmyo-3::mCherry*) at 5 ng/µL, pCFJ90 (*Pmyo-2::mCherry*) at 2.5 ng/µL, pMA122 (*Phsp16.41::peel-1*) at 10 ng/µL, and carrier DNA Litmus 38i to a final concentration of 100 ng/µL DNA.

Extrachromosomal arrays were integrated into the genome by exposure to 4000 rads of gamma irradiation. Irradiated young adult hermaphrodites were transferred to 10 cm OP50 plates (5 worms/plate) and grown to starvation. The plates were chunked and grown to starvation twice more to enrich for stably expressing lines. When nearly starved, 8 animals per plate were picked to individual plates. The progeny were then screened for 100% stable transmission, indicating integration into the genome. Integration was confirmed by mapping the transgene to a chromosome.

### Aldicarb assays

35 mm aldicarb assay plates were poured with NGM supplemented with 1 mM aldicarb. The plates were seeded with 5 uL OP50 and dried at room temperature overnight. Animals were picked onto the OP50 lawn to begin the assay (time 0) and then kept at room temperature. Every 15 minutes, animals were scored for paralysis by lightly touching the nose of the animal with an eyebrow hair. Animals were scored as paralyzed if the worm displayed no locomotor response to three nose touches and had no pharyngeal pumping. Animals that left the OP50 lawn were picked back onto the food.

### Statistical analysis

P values were determined using GraphPad Prism 5. Normally distributed data sets were analyzed with a one-way ANOVA and Bonferroni’s *post hoc* test when group size was unequal, or with Tukey’s *post hoc* test when group size was equal. Data sets with non-normal distribution (using the Shapiro-Wilk normality test) were analyzed with a Kruskal-Wallis test and Dunn’s *post hoc* test. Data sets with multiple independent variables were analyzed by two-way ANOVA and Bonferroni’s *post hoc* test.

### Reagent and data availability

Strains and plasmids are listed in Tables S1 and S2 and are available upon request. Primers are listed in Table S3. The authors state that all data necessary for confirming the conclusions presented in the article are represented fully within the article and Supplemental Material.

## Results

### KSR-1 regulates locomotion downstream of or in parallel to Gq

In *C. elegans,* the heterotrimeric G protein Gq regulates synaptic vesicle release (Hu *et al.* 2015). Gq is a key regulator of neuromuscular activity, as loss-of-function mutants in *egl-30* are nearly paralyzed (Brundage *et al.* 1996) whereas the gain-of-function mutant *egl-30(tg26)* has hyperactive locomotion with an exaggerated loopy waveform (Doi and Iwasaki 2002; Bastiani *et al.* 2003) (Figure 1, A, C, and D). To identify pathways required for Gq signaling, we performed a forward genetic screen in *C. elegans* for suppressors of the activated Gq mutant *egl-30*(*tg26)*. This screen has been used to identify new components of the Gq signal transduction pathway and proteins important for dense-core vesicle function (Williams *et al.* 2007; Ailion *et al.* 2014; Topalidou *et al.* 2016; Topalidou *et al.* 2017a; b). Two previously uncharacterized suppressors identified in this screen, *ox314* and *yak10*, showed similar suppression of the loopy waveform and hyperactivity of *egl-30(tg26)* animals (Figure 1D). When crossed away from the *egl-30(tg26*) background, both mutants moved with wild-type waveform (Figure 1D), but at a slightly slower rate. We mapped the *ox314* allele near the center of the X chromosome (see Materials and Methods), and a complementation test showed that *ox314* and *yak10* are mutations in the same gene since they fail to complement in an *egl-30(tg26)* background.

We used whole genome sequencing to identify *ox314* and *yak10* as nonsense mutations in *ksr-1* (Figure 1B, see Materials and Methods). KSR-1 is a scaffold protein that facilitates the localization and interactions required for the Ras-mitogen activated protein kinase (MAPK) cascade consisting of Raf, MEK, and ERK (Kornfeld *et al.* 1995b; Sundaram and Han 1995; Nguyen *et al.* 2002; Zhang *et al.* 2013). The deletion allele *ksr-1(ok786)* also suppressed the loopy waveform of the activated Gq mutant identically to *ox314* and *yak10*. These results suggest that KSR-1 activity is required for regulation of locomotion rate and waveform by Gq.

### The ERK MAPK cascade acts to promote Gq signaling

Because the loss of the MAPK scaffold *ksr-1* suppresses the activated Gq mutant *egl-30(tg26)*, we asked whether other components of the Ras-ERK pathway would also suppress. Since the core components of the Ras-ERK pathway are required for viability, we built double mutants of activated Gq with reduction-of-function mutations in genes at each step of the ERK cascade. Mutations in Raf (*lin-45(sy96)*), MEK (*mek-2(n1989)*, *mek-2(ku114)*), and ERK (*mpk-1(ga117)*, *mpk-1(oz140)*) all suppressed the loopy waveform of activated Gq animals similarly to *ksr-1(ok786)* (Figure 2, A and B; Figure S1, A and B). However, mutations in Ras (*let-60(n2021)*) and the upstream pathway activators EGF (*lin-3(e1417)*) and the EGF receptor (*let-23(sy12)*) did not suppress activated Gq (Figure 2C). Because *let-60* is required for viability, most *let-60* alleles including *n2021* are partial loss-of-function (Beitel *et al.* 1990). We also analyzed the dominant negative D119N allele *let-60(sy93)* that disrupts Ras binding to guanine nucleotides and thus prevents Ras activation (Han and Sternberg 1991). We found that *let-60(sy93)* also did not suppress the loopy waveform of activated Gq (Figure 2D). Additionally, we expressed the dominant negative *let-60* mutation S17N specifically in neurons and did not observe suppression of the loopy waveform of activated Gq (Figure S2D). These results support the possibility that ERK activation in this pathway occurs through a Ras-independent mechanism.

**Figure 2.**
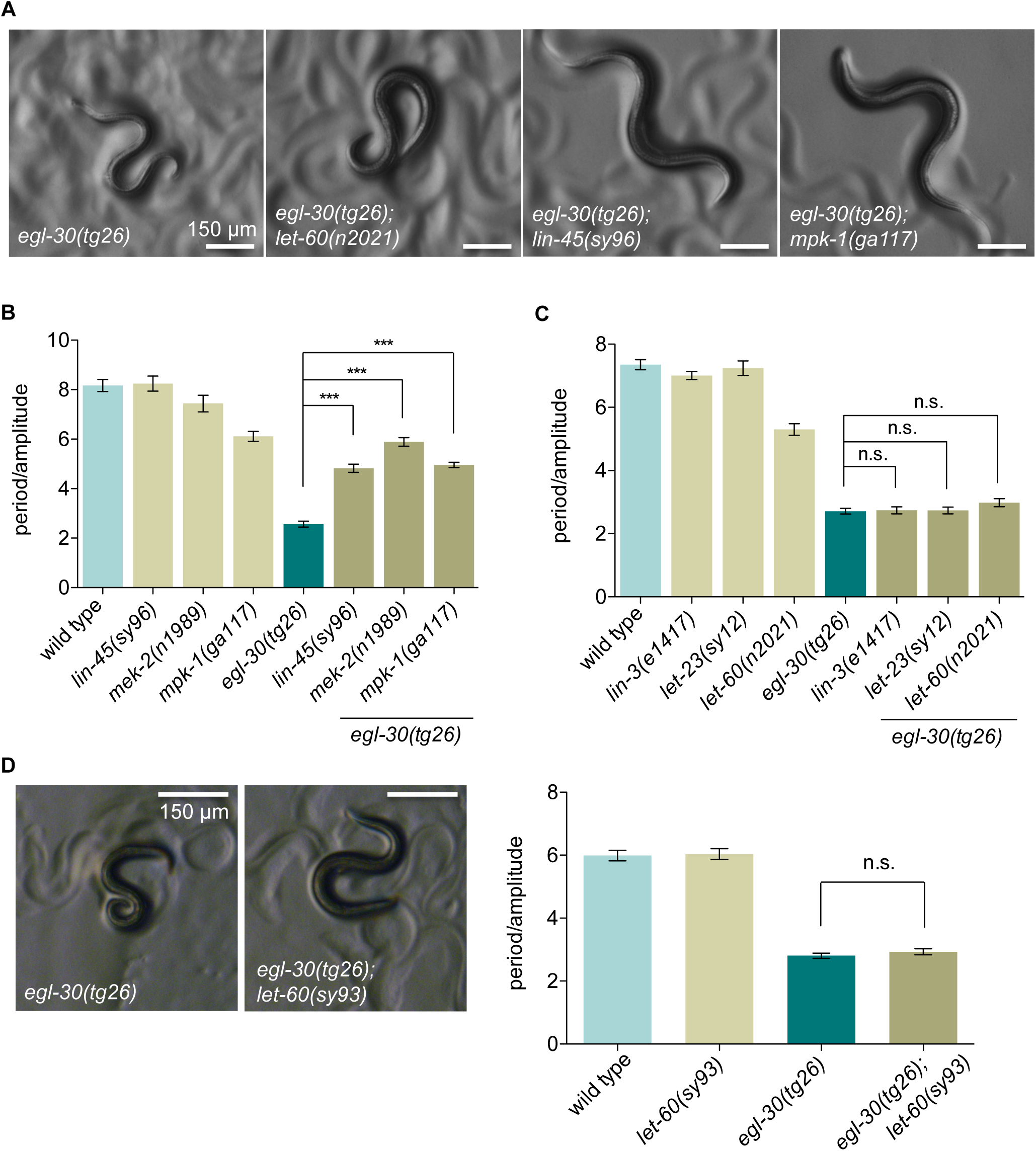
Mutations in the ERK MAPK pathway suppress activated Gq.(A) Mutations in the MAPKKK *lin-45*/Raf and the MAPK *mpk-1*/ERK suppress the coiled posture of *egl-30* (*tg26*) worms. Partial loss of Ras *let-60* activity does not suppress activated Gq posture. (B)Waveform quantification of ERK pathway mutants. Mutations in *lin-45*, *mek-2*, and *mpk-1* suppress activated Gq *egl-30*(*tg26*) similar to *ksr-1* mutations. *N ≥* 12, *** *P* < 0.001, one-way ANOVA with Bonferroni’s *post hoc* test. (C) Signaling components upstream of Raf do not suppress activated Gq. Mutations in the EGF ligand (*lin-3*) or the EGFR (*let-23*) do not affect the exaggerated waveform of *egl-30*(*tg26*) animals. The Ras partial loss-of-function mutation *let-60(n2021)* does not suppress activated Gq waveform. *N ≥* 12, *** *P* < 0.001, one-way ANOVA with Bonferroni’s *post hoc* test. (D)The *let-60(sy93)* dominant negative mutation in Ras does not suppress activated Gq waveform. *N ≥* 13, n.s., not significant, one-way ANOVA with Bonferroni’s *post hoc* test.

Because partial loss-of-function mutations in the ERK MAPK pathway genes downstream of Ras showed clear suppression of activated Gq, we were surprised to find that partial loss-of-function mutations in Ras did not suppress. If LET-60/Ras is indeed not required, Gq might instead activate the ERK pathway via other Ras-subfamily proteins. To test this possibility, we made double mutants of activated Gq with putative null alleles of R-Ras/*ras-1*, M-Ras/*ras-2*, and Rap1/*rap-1* and found that they also did not suppress activated Gq (Figure S2A). To further investigate whether this pathway acts independently of Ras, we tested mutations in GEFs that activate Ras. The temperature-sensitive RasGEF mutant *sos-1(cs41)* did not suppress activated Gq when shifted to the non-permissive temperature (Figure S2B). Additionally, a null mutation in the neuronal RasGEF *rgef-1* also did not suppress activated Gq (Figure S2C). In genetic screens for vulval induction mutants, additional factors such as the PP2A subunit *sur-6* (Sieburth *et al.* 1999) and ion transporter *sur-7* (Yoder *et al.* 2004) were identified as positive regulators of Ras-ERK activity. However, the *sur-6(sv30)* and *sur-7(ku119)* mutations did not suppress activated Gq locomotion (data not shown). These data suggest either that only a low level of Ras activity is needed to properly activate ERK signaling downstream of Gq, or that ERK signaling acts independently of LET-60/Ras and other known *C. elegans* Ras family proteins to regulate locomotion downstream of Gq.

### KSR-1 and the ERK MAPK cascade modulate Rho signaling

Three classes of suppressor mutations were isolated in our forward genetic screen of activated Gq, as characterized by their molecular roles and unique suppression phenotypes (Topalidou *et al.* 2017a; b). We grouped together a class of suppressor mutations including *ox314, yak10*, and the RhoGEF Trio (*unc-73* in *C. elegans*) by their characteristic strong suppression of the loopy waveform of activated Gq (Topalidou *et al.* 2017a; b), suggesting that *ksr-1* might act in the same pathway as *unc-73*.

We have shown that Gq regulates locomotion via the small GTPase Rho (RHO-1 in *C. elegans*) (Topalidou *et al.* 2017a). Transgenic expression of an activated RHO-1 mutant (G14V) in acetylcholine neurons (here called “*rho-1(gf)*”) causes worms to have a loopy waveform and impaired locomotion (McMullan *et al.* 2006) (Figure 3A). To examine whether *ksr-1* acts in the Gq-Rho pathway we tested whether mutations in *ksr-1* suppress the phenotypes of *rho-1(gf)* worms. We found that the *ksr-1* alleles *ok786, ox314,* and *yak10* all suppressed the loopy waveform of *rho-1(gf)* worms (Figure 3, A, B, and D). Because *rho-1(gf)* worms have a slow locomotion rate and loopy waveform, these mutants do not efficiently travel long distances. We used radial locomotion assays (see Materials and Methods) to quantify the locomotion phenotype of *rho-1(gf)* worms. *rho-1(gf) ksr-1* double mutants had an increased radial distance traveled compared to *rho-1(gf)* alone (Figure 3B). These data suggest that KSR-1 acts downstream of or in parallel to the Gq-Rho pathway to regulate locomotion.

**Figure 3.**
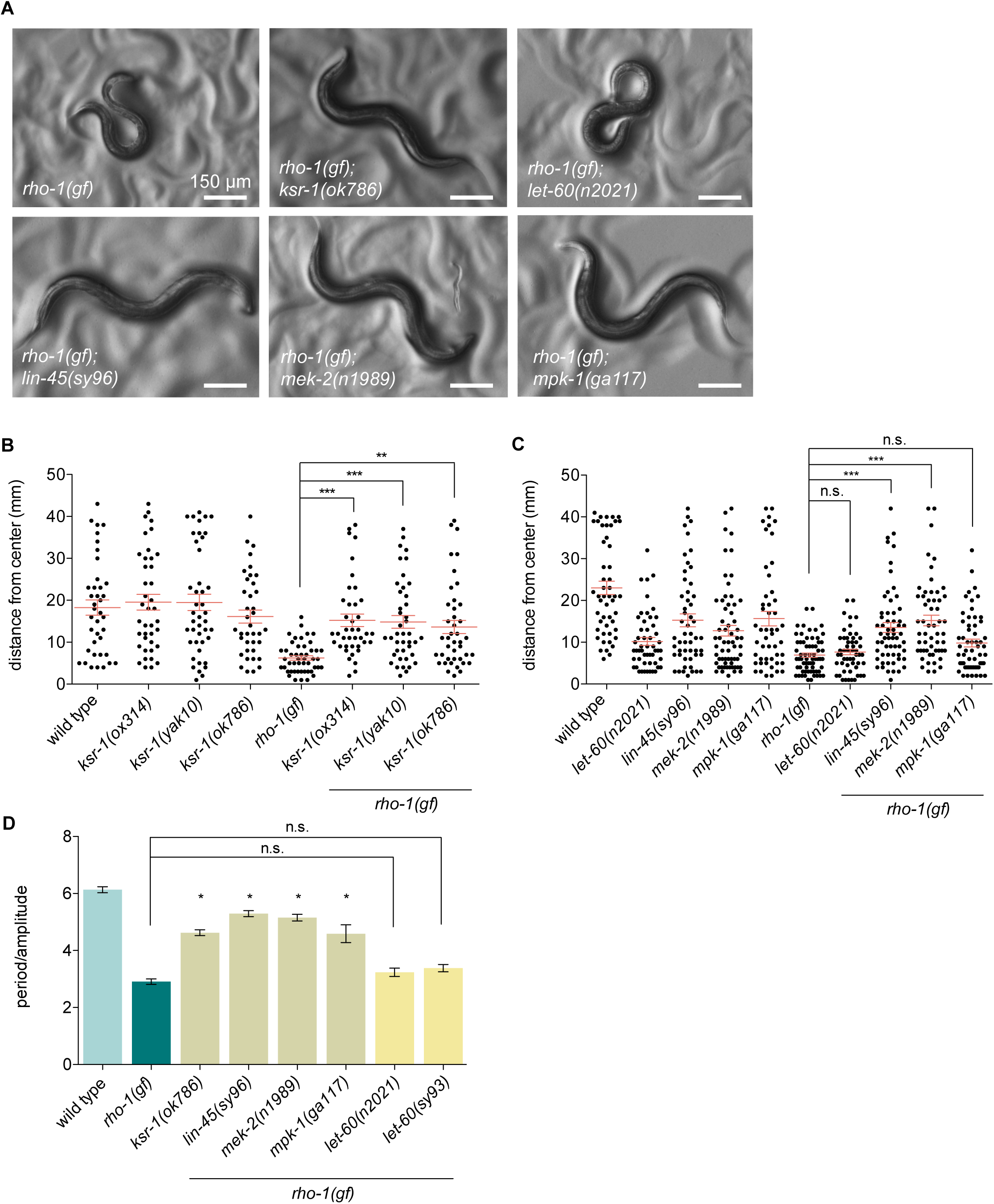
Mutations in *ksr-1* and the ERK pathway suppress activated Rho. (A) Mutations in ERK pathway genes *ksr-1*, *lin-45*, *mek-2*, and *mpk-1* visibly suppress the coiled posture of animals expressing an activated Rho mutant (G14V) in acetylcholine neurons (*nzIs29[Punc-17::rho-1(gf)]*). Reduction of LET-60/Ras activity does not suppress activated Rho. (B)The *ksr-1*(*ok786*)*, ksr-1*(*ox314*), and *ksr-1*(*yak10*) mutations suppress the locomotion of activated Rho animals as shown by radial locomotion assays. *N ≥* 38, *** *P* < 0.001, ** *P <* 0.01, Kruskal-Wallis test with Dunn’s *post hoc* test. (C) Mutations in *lin-45* and *mek-2* suppress the locomotion defect of activated Rho animals as shown by radial locomotion assays. The *let-60*(*n2021*) mutation does not significantly suppress activated Rho locomotion. *N ≥* 50, *** *P* < 0.001, Kruskal-Wallis test with Dunn’s *post hoc* test. (D) Waveform quantification of ERK pathway mutants. Mutations in *ksr-1*, *lin-45*, *mek-2*, and *mpk-1* show similar suppression of activated Rho, but the Ras reduction-of-function mutant *let-60(n2021)* or dominant negative *let-60(sy93)* do not suppress activated Rho. N ≥ 14, * P < 0.05, n.s., not significant, one-way ANOVA with Bonferroni’s *post hoc* test, compared to *rho-1(gf)*.

Since *ksr-1* mutants suppress the exaggerated waveform of both activated Gq and activated Rho animals, we expected that loss of other ERK pathway components would also suppress activated Rho. We made double mutants of activated Rho (*rho-1(gf)*) with reduction-of-function alleles of the Ras-ERK pathway and found that mutations in Raf, MEK, and ERK suppressed the *rho-1(gf)* loopy waveform phenotype (Figure 3, A, C, and D). However, the *let-60(n2021)* and *let-60(sy93)* Ras mutations did not suppress the loopy waveform of *rho-1(gf)* worms (Figure 3, A, C, and D). These data suggest that the ERK pathway acts downstream of or in parallel to the Gq-Rho pathway to regulate locomotion, possibly in a Ras-independent manner.

### The ERK MAPK cascade acts in acetylcholine neurons to control locomotion

Members of the ERK pathway are expressed in neurons in *C. elegans* (Dent and Han 1998; Hunt-Newbury *et al.* 2007), and Gq and Rho act in acetylcholine neurons to promote synaptic release and regulate locomotion (Lackner *et al.* 1999; McMullan *et al.* 2006). To determine whether the ERK pathway also acts in neurons to modulate Gq signaling, we expressed the *ksr-1* cDNA under promoters driving expression in specific types of neurons. Single-copy transgenic expression of *ksr-1* under an acetylcholine neuron promoter (*Punc-17*) or under a head acetylcholine neuron promoter (*Punc-17H)* fully reversed the *ksr-1* suppression of the loopy waveform of activated Gq worms (Figure 4). *ksr-1* expression in ventral cord acetylcholine motor neurons (*Punc-17β*) or GABA neurons (*Punc-47)* did not significantly reverse the *ksr-1* suppression of activated Gq (Figure 4). This suggests that ERK signaling primarily functions in the acetylcholine interneurons or motor neurons of the head to modulate Gq-dependent locomotion.

**Figure 4.**
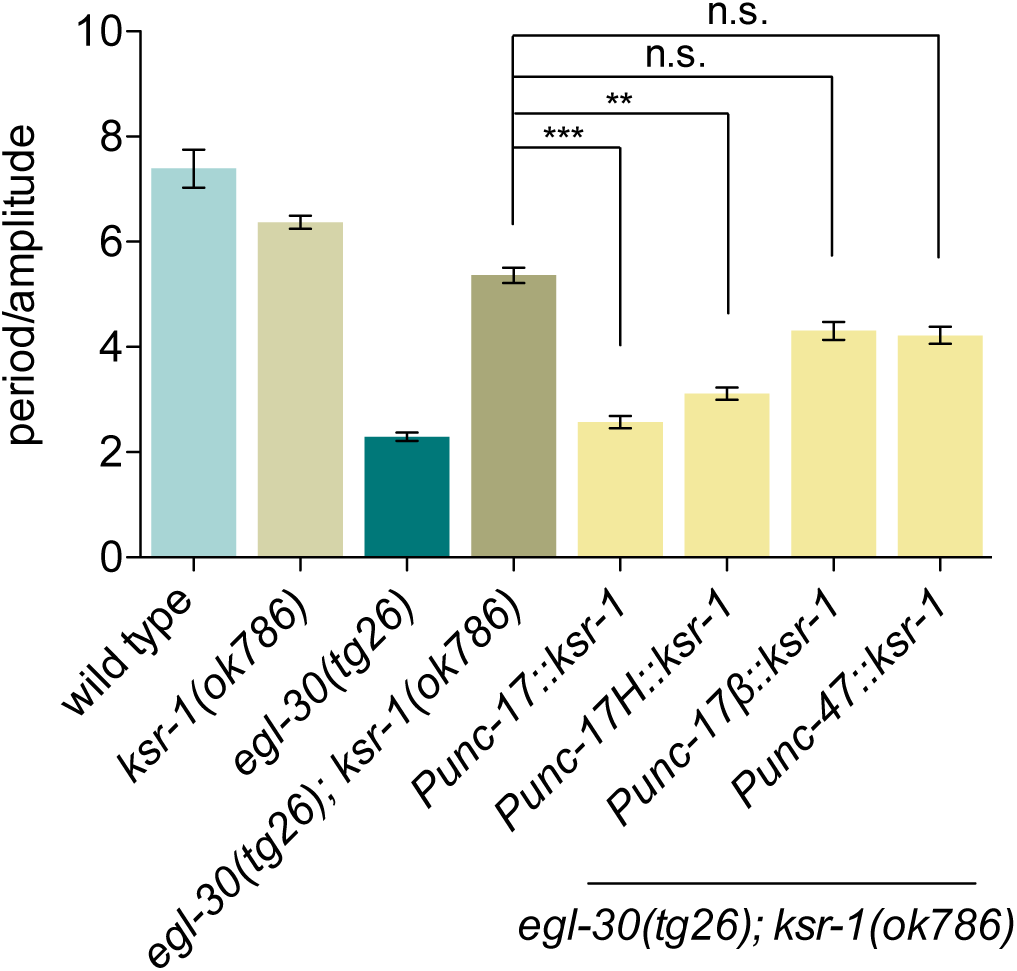
KSR-1 acts in head acetylcholine neurons to modulate Gq signaling. Single-copy expression of the *ksr-1* cDNA exclusively in acetylcholine neurons (*Punc-17*, *yakSi26* transgene) or head acetylcholine neurons (*Punc-17H, yakSi27* transgene) is sufficient to reverse the *ksr-1* suppression of the loopy waveform of activated Gq animals. *ksr-1* expression in ventral cord acetylcholine neurons (*Punc-17β*, *yakSi28* transgene) or GABA neurons (*Punc-47*, *yakSi29* transgene) does not reverse the *ksr-1* suppression of the activated Gq exaggerated waveform. *N ≥* 12, *** *P* < 0.001, ** *P* < 0.01, n.s., not significant, Kruskal-Wallis test with Dunn’s *post hoc* test.

We have shown that the ERK pathway is necessary for Gq-dependent effects on locomotion. To determine whether ERK signaling is sufficient to modulate locomotion, we expressed an activated form of *lin-45*/Raf specifically in acetylcholine neurons. Raf kinase activity is regulated via conserved phosphorylation events, and phosphomimetic mutations T626E/T629D in the kinase activation loop of *lin-45* are sufficient to confer constitutive Raf activity (Chong *et al.* 2001). We found that expression of activated Raf in acetylcholine neurons (*Punc-17*) causes a loopy waveform similar to activated Gq and Rho mutants and similar limited dispersal in radial locomotion assays (Figure 5, A and B; Figure S3, A and B). Additionally, expression of activated Raf or activated Gq in head acetylcholine neurons also caused a loopy waveform (Figure S3, A-C).

**Figure 5.**
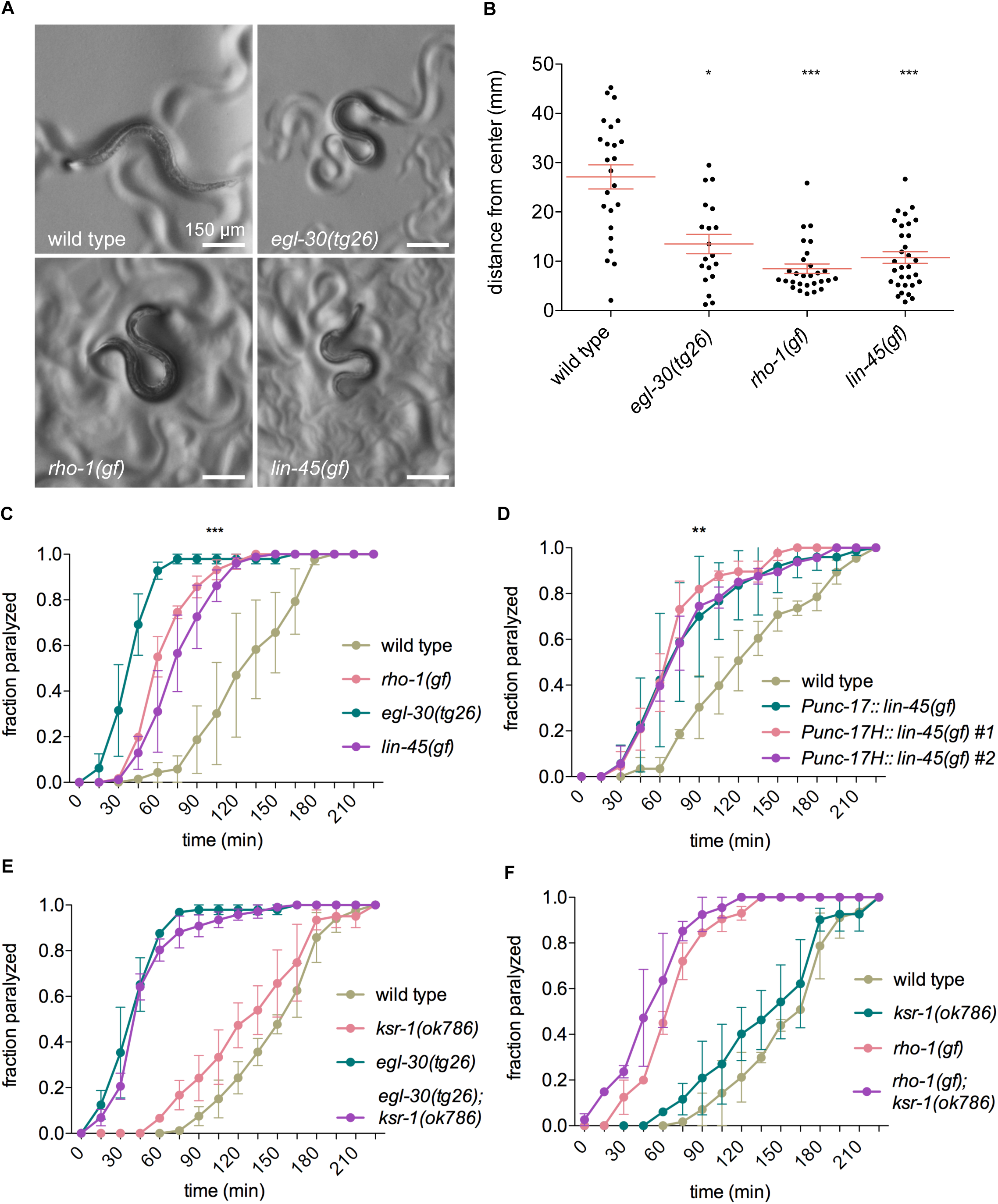
Increased Raf activity in acetylcholine neurons is sufficient to regulate locomotion and increase acetylcholine release. (A) Transgenic lines expressing an activated form of *lin-45*/Raf (T626E/T629D) in acetylcholine neurons (*yakIs34[Punc-17::lin-45(gf)]*) have exaggerated body bends and coiled posture similar to the activated Gq mutant *egl-30*(*tg26*) and to animals expressing activated Rho in acetylcholine neurons (*nzIs29[Punc-17::rho-1(gf)]*). The wild type and *egl-30(tg26)* photos are the same as shown in Figure 1A, and the *rho-1(gf)* photo is the same as the one in Figure 3A. (B) Expression of activated Rho (*nzIs29[Punc-17::rho-1(gf)]*) or Raf (*yakIs34[Punc-17::lin-45(gf)]*) in acetylcholine neurons impairs coordinated locomotion similarly to activated Gq (*egl-30*(*tg26*)). *N ≥* 19, *\* P* < 0.05, *** *P* < 0.001, Kruskal-Wallis test with Dunn’s *post hoc* test. (C) Animals expressing activated Gq, Rho, or Raf are hypersensitive to the acetylcholinesterase inhibitor aldicarb. Activated Gq (*egl-30*(*tg26*)), activated Rho expressed in acetylcholine neurons (*nzIs29[Punc-17::rho-1(gf)]*), and activated Raf expressed in acetylcholine neurons *yakIs34[Punc-17::lin-45(gf)]*) become paralyzed significantly faster than wild type animals when exposed to 1 mM aldicarb. All strains are significantly different from wild type at t = 60, 75, 90, and 105 minutes. *N ≥* 61, *** *P* < 0.001, two-way ANOVA with Bonferroni’s *post hoc* test. (D) Animals expressing activated Raf in head acetylcholine neurons are hypersensitive to aldicarb. The *yakIs34[Punc-17::lin-45(gf)]* integrated array expressing activated Raf in all acetylcholine neurons and the *yakEx154[Punc-17H::lin-45(gf)]* (#1) and *yakEx168[Punc-17H::lin-45(gf)]* (#2) extrachromosomal arrays expressing activated Raf in head acetylcholine neurons show similar hypersensitivity to aldicarb. All strains are significantly different from wild type at t = 60, 75, 90, and 105 minutes. N ≥ 47, ** P < 0.01, two-way ANOVA with Bonferroni’s *post hoc* test. (E) *ksr-1* is not required for the aldicarb hypersensitivity of activated Gq. Paralysis of *ksr-1*(*ok786*) animals on 1 mM aldicarb is not significantly different from wild type. The *ksr-1* deletion *ok786* does not suppress the aldicarb hypersensitivity of activated Gq (*egl-30*(*tg26*)). *N ≥* 53. (F) *ksr-1* is not required for the aldicarb hypersensitivity of activated Rho. Paralysis of *ksr-1(ok786)* animals on 1 mM aldicarb is not significantly different from wild type. The *ksr-1* deletion *ok786* does not suppress the aldicarb hypersensitivity of the activated Rho mutant *nzIs29[Punc-17::rho-1(gf)]*. N ≥ 41.

Gq and Rho promote acetylcholine release at the worm neuromuscular junction (Lackner *et al.* 1999; Miller *et al.* 1999; McMullan *et al.* 2006). To determine if Raf activation affects acetylcholine release at the neuromuscular junction, we assayed for sensitivity to the acetylcholinesterase inhibitor aldicarb. Mutants with reduced acetylcholine secretion are resistant to aldicarb, whereas mutants with increased acetylcholine secretion are hypersensitive to aldicarb (Mahoney *et al.* 2006). Activated Gq and Rho mutants have increased rates of paralysis when exposed to aldicarb (Lackner *et al.* 1999; McMullan *et al.* 2006). We found that expression of activated Raf in all acetylcholine neurons or in head acetylcholine neurons also led to aldicarb hypersensitivity, similar to activated Gq and Rho mutants (Figure 5, C and D). However, we found that a *ksr-1* mutation does not suppress the aldicarb hypersensitivity of activated Gq or Rho mutants, and the *ksr-1* mutant on its own has similar aldicarb sensitivity to wild type (Figure 5, E and F). These results suggest that the ERK pathway is not required for synaptic transmission, but is sufficient to stimulate synaptic transmission when constitutively activated. Furthermore, these data suggest that ERK pathway activation in head acetylcholine neurons is sufficient to promote acetylcholine release by motor neurons at the neuromuscular junction.

## Discussion

In this study we identified KSR-1 and the ERK MAPK cascade as modulators of Gq-Rho signaling. We found that ERK signaling acts in head acetylcholine neurons (interneurons or motor neurons of the head) to modulate Gq-dependent locomotion behavior, especially the waveform of the animal. Our data support the model that Gq-Rho activation of the ERK pathway may be independent of its canonical regulator, the small GTPase Ras/LET-60 (Figure 6).

**Figure 6.**
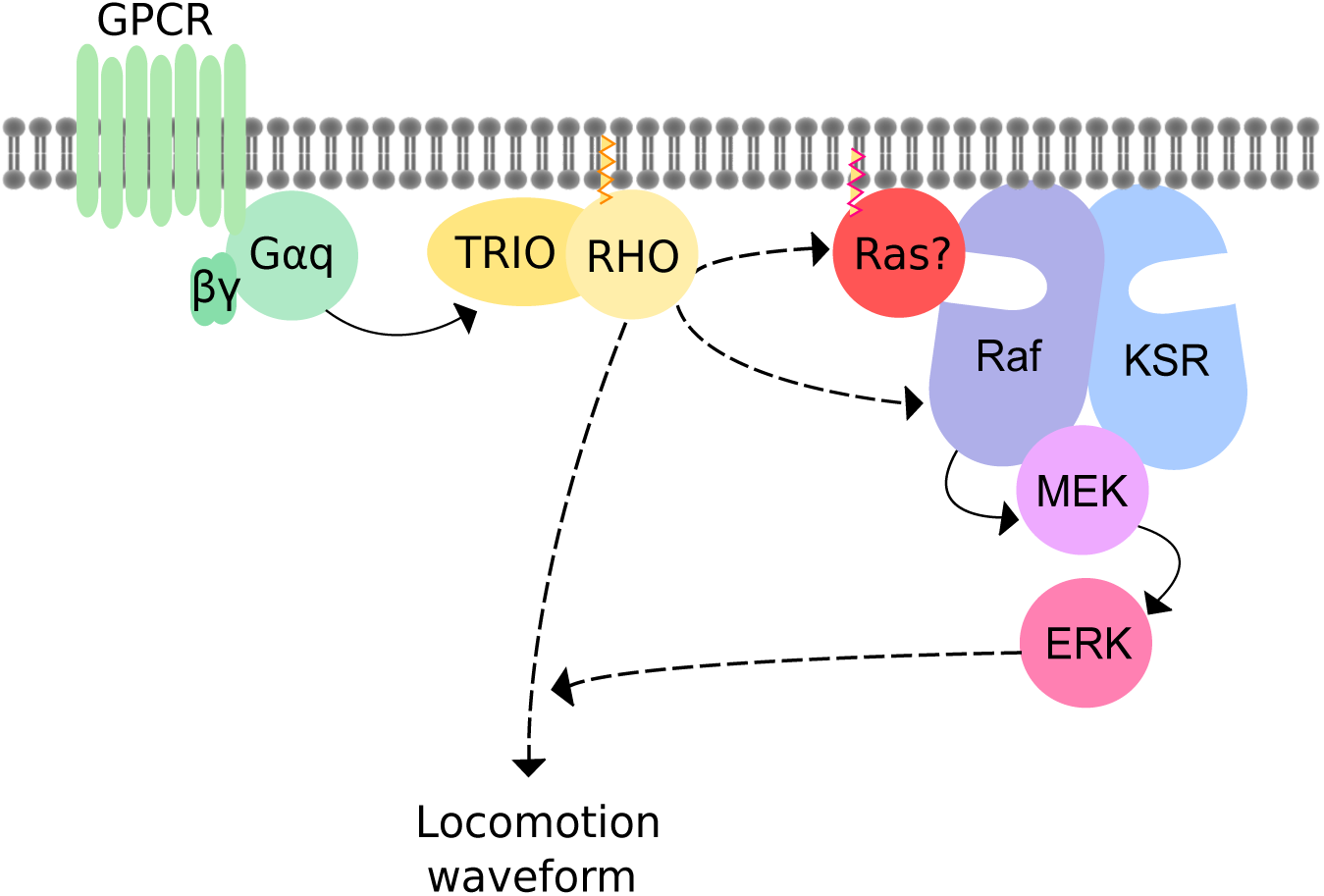
Model for Gq-Rho-ERK signaling. Gq directly activates Trio and Rho (solid arrow) (Lutz et al. 2007). The core ERK cascade acts either downstream of Rho or in parallel (dashed arrows) to modulate locomotion behavior. ERK activation occurs independently of the extracellular growth factor LIN-3, its receptor LET-23, or Ras/LET-60. It is possible that Raf/LIN-45 is activated by a Ras family member other than the canonical worm Ras/LET-60. Our model suggests that Gq-Rho signaling regulates the worm waveform by acting cell autonomously in head acetylcholine neurons in an ERK-dependent pathway. By contrast, we propose that Gq-Rho signaling regulates synaptic transmission at the neuromuscular junction by acting in an ERK-independent pathway, possibly in the ventral cord motorneurons themselves.

Gq regulates multiple aspects of *C. elegans* locomotion behavior, including the rate of locomotion and waveform. Previous work has shown that these behaviors are at least partially genetically separable, with different pathways downstream of Gq contributing differentially to these behaviors and with Gq likely acting in several classes of neurons including both head acetylcholine neurons and ventral cord acetylcholine motor neurons (Lackner *et al.* 1999; Topalidou *et al.* 2017a). Here we focused on Gq regulation of the waveform, which is strongly dependent on Rho and ERK. The most parsimonious model supported by our findings is that Gq, Rho, and the ERK pathway function cell autonomously in the head acetylcholine neurons to regulate the waveform (Figure 6). Other components of the Gq-Rho pathway have been previously shown to act in head acetylcholine neurons to regulate the waveform (Topalidou *et al.* 2017a; b), and here we show that KSR-1 is required in head acetylcholine neurons to modulate Gq-dependent effects on the waveform. Additionally, expression of activated LIN-45/Raf in head acetylcholine neurons is also sufficient to regulate the waveform and cause loopy locomotion. These data suggest that ERK pathway activity in head acetylcholine neurons is both necessary and sufficient to regulate worm waveform.

Though the ERK pathway is required for Gq and Rho-dependent regulation of the waveform, it is not required for the aldicarb hypersensitivity of activated Gq and Rho mutants. This indicates that Gq and Rho regulation of acetylcholine release at the neuromuscular junction is at least partially independent of Gq and Rho regulation of the locomotion waveform. The simplest model is that Gq and Rho signaling regulates the waveform by acting in head acetylcholine neurons through an ERK-dependent pathway, but Gq and Rho signaling regulates synaptic transmission at the neuromuscular junction through an ERK-independent pathway. Overexpression of an activated Gq mutant in ventral cord acetylcholine motor neurons is sufficient to cause an aldicarb hypersensitive phenotype (Lackner *et al.* 1999), suggesting that Gq may act cell autonomously in the ventral cord motor neurons to promote acetylcholine release by these neurons. But as with activated Raf, it is also possible that Gq activity in head acetylcholine neurons may contribute to acetylcholine release by downstream ventral cord motor neurons.

The ERK signaling cascade has been well-studied for its regulation of cellular proliferation and differentiation (Sun *et al.* 2015), but also plays important roles in mature neurons and has been associated with synaptic plasticity and memory (Impey *et al.* 1999). In addition to activating transcription, ERKs regulate synaptic plasticity both presynaptically and postsynaptically by phosphorylating relevant substrates. ERKs phosphorylate the presynaptic proteins synapsin I and Munc18-1, and postsynaptic proteins such as scaffolds, K_v_4.2 potassium channels, and Group I metabotropic glutamate receptors (Jovanovic *et al.* 2000; Thomas and Huganir 2004; Sweatt 2004; Kushner *et al.* 2005; Boggio *et al.* 2007; Vara *et al.* 2009; Mao and Wang 2016a; Schmitz *et al.* 2016). Our findings suggest that the ERK pathway controls Gq-dependent locomotion in *C. elegans* and that activated ERK promotes synaptic transmission.

In contrast to many developmental and neuronal roles of Ras-dependent ERK signaling in *C. elegans,* our data suggest that Raf-MEK-ERK signaling may modulate Gq-Rho output independently of Ras/LET-60. One caveat to the conclusion that the ERK pathway acts independently of Ras to control Gq-dependent locomotion is that the alleles of *let-60*/Ras tested here are not null (Han and Sternberg 1990, 1991; Han *et al.* 1990; Beitel *et al.* 1990). However, other than *mpk-1(ga117)*, the alleles we used of *lin-45*, *mek-2* and *mpk-1* are also not null (Han *et al.* 1993; Kornfeld *et al.* 1995a; Wu *et al.* 1995; Lackner and Kim 1998; Hsu *et al.* 2002), yet were able to suppress an activated Gq mutant. Furthermore, the *let-*60/Ras alleles used here have stronger phenotypes in vulval development than the two weak *mek-2*/MEK alleles we used (*n1989* and *ku114*) (Beitel *et al.* 1990; Kornfeld *et al.* 1995a; Wu *et al.* 1995) and the *let-60(n2021)*/Ras mutant has comparable phenotypes to the *lin-45(sy96)*/Raf and *mpk-1(ga117*)/ERK alleles for chemotaxis to odorants and the regulation of foraging behavior (Hirotsu *et al.* 2000; Hamakawa *et al.* 2015). Thus, either LET-60/Ras is not required to modulate Gq signaling, or ERK signaling in the locomotor circuit requires a lower threshold of Ras activity and the *let-60* mutants we used have sufficient levels of active Ras to modulate Gq signaling, but not enough for vulval induction or other behaviors.

If Ras is not required, how instead might KSR-Raf-MEK-ERK signaling be activated? Normally, active Ras recruits Raf to the plasma membrane but it has been shown that artificial recruitment of Raf to the plasma membrane in the absence of Ras is sufficient to activate Raf signaling (Stokoe *et al.* 1994; Marais *et al.* 1995). Thus, another possible way to activate the ERK pathway would be for Raf to be recruited to the membrane by proteins other than Ras. The most obvious candidates for recruiting Raf are other members of the Ras family such as Rap1 or R-Ras that have a conserved effector-binding domain (Reiner and Lundquist 2016); it has been reported that Rap1 can mediate Ras-independent activation of Raf downstream of a Gq-coupled receptor (Guo *et al.* 2001). However, mutations in the worm orthologs of Rap1/RAP-1 and R-Ras/RAS-1 did not suppress the activated Gq mutant, indicating that these Ras-like proteins are not required, at least individually, to activate Raf in this pathway. It is possible that more than one of these Ras family members function redundantly to activate Raf, or that Raf is activated independently of Ras family proteins. There have been other reported cases of Ras-independent activation of the Raf-MEK-ERK pathway (Robbins *et al.* 1992; Honda *et al.* 1994; van Biesen *et al.* 1996; Ueda *et al.* 1996; Drosten *et al.* 2014), some of which involve G protein signaling and protein kinase C, though the precise mechanisms involved are unclear.

Gq, Rho, and ERK are also required for the *C. elegans* behavioral and immune response to infection by the bacterium *M. nematophilum* (Nicholas and Hodgkin 2004; McMullan *et al.* 2012). Interestingly, LET-60/Ras was reported to not be required for the immune response (Nicholas and Hodgkin 2004) or only partially required (McMullan *et al.* 2012). Additionally, LET-60/Ras was not required for the increased sensitivity to aldicarb caused by bacterial infection (McMullan *et al.* 2012), at least as assayed using the *let-60(n2021*) allele. Thus, the same neuronal Ras-independent ERK pathway we describe here may also modulate Gq signaling in the neuronal response to bacterial infection and possibly the innate immune response as well.

Our epistasis analysis suggests that ERK signaling acts genetically downstream of or in parallel to Gq-Rho signaling. How might Gq-Rho signaling lead to ERK activation? ERK activation could occur via a linear pathway downstream of Gq and Rho, or ERK could signal in parallel and converge downstream of Rho to affect neuronal activity (Figure 6). There is precedence for Gq activating ERK via a linear pathway. In the pharyngeal muscle of *C. elegans*, ERK activity is increased by Gq-dependent signaling through protein kinase C (You *et al.* 2006). In the AWC olfactory neuron, ERK activity is increased downstream of Gq signaling via the RasGEF RGEF-1 (Chen *et al.* 2011; Uozumi *et al.* 2012). Protein kinase C and RGEF-1 are both activated by DAG, probably produced by the canonical Gq-PLCβ pathway. By contrast, we found that ERK signaling regulates locomotion by modulating the output of the Gq-Rho pathway that acts in parallel to Gq-PLCβ and does not depend on RGEF-1.

How might the ERK pathway modulate neuronal activity downstream of Gq-Rho signaling? Rho regulates neuronal activity and synaptic release through several mechanisms in *C. elegans* neurons, any of which could be targets of ERK signaling. One possible ERK effector is the NCA/NALCN cation channel that acts genetically downstream of Gq-Rho to regulate locomotion rate and waveform (Topalidou *et al.* 2017a; b). Though ERK has not been directly connected to NCA/NALCN, mammalian ERK regulates neuronal excitability by directly phosphorylating voltage-gated sodium and potassium channels (Schrader *et al.* 2006; Stamboulian *et al.* 2010) and by regulating channel expression (Yang *et al.* 2015). Given the many possible ways that ERK may regulate excitability or synaptic transmission, *C. elegans* genetics is well suited to determine relevant effectors of Gq-Rho-ERK signaling.

## Acknowledgements

We thank Brooke Jarvie for the isolation of *ksr-1(yak10)*; Stephen Nurrish for the activated Rho worm strain; Jordan and Jill Hoyt for help with whole-genome sequence analysis; Laura Taylor for help with irradiation of extrachromosomal arrays; Dana Miller for use of her Nikon SMZ18 microscope and camera. Some strains were provided by the CGC, which is funded by the NIH Office of Research Infrastructure Programs (P40 OD010440). This work was supported by an Ellison Medical Foundation New Scholar Award and by NIH grants R00 MH082109 and R56 NS100843 to M.A.

## Figure Legends

**Figure S1.**
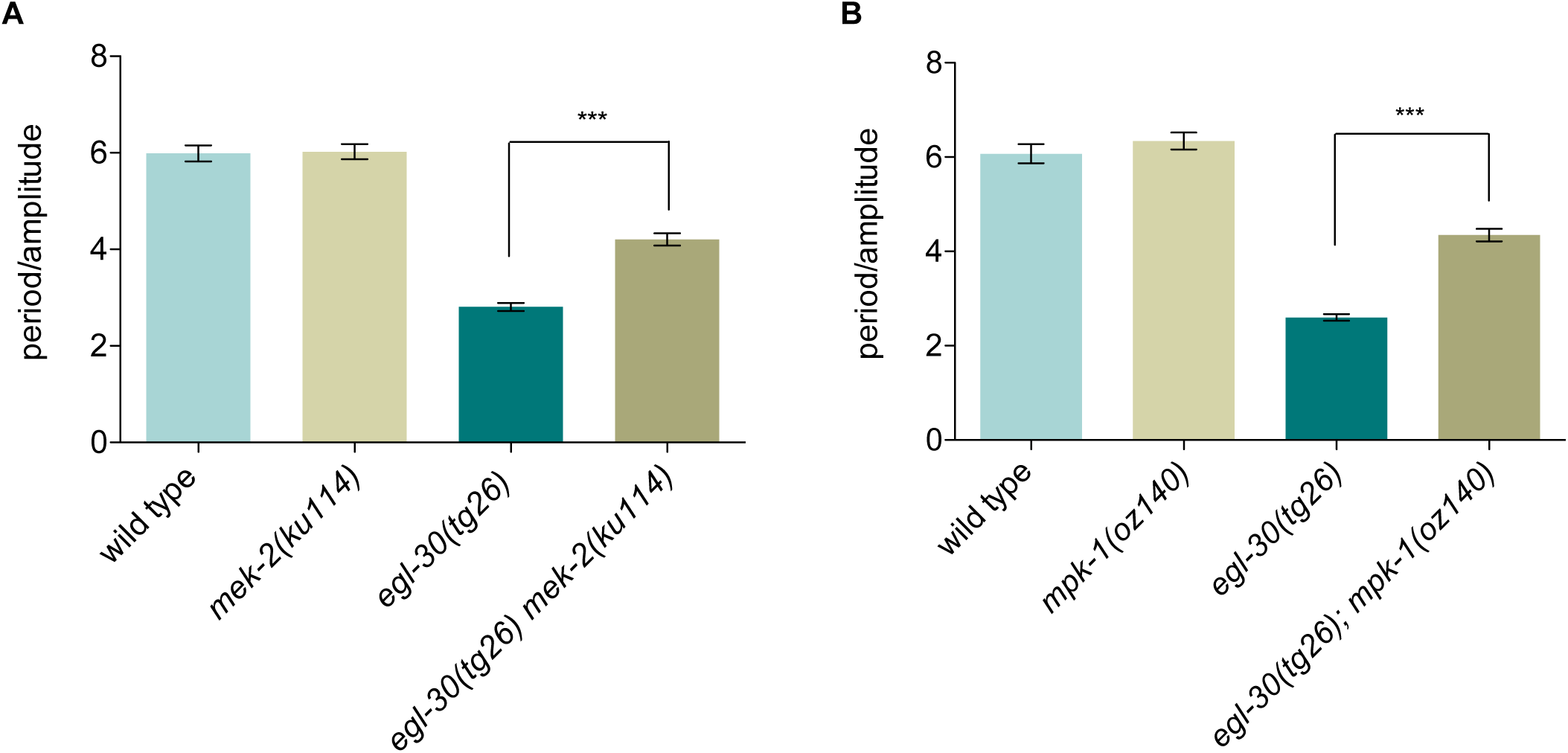
Additional mutations in *mek-2* and *mpk-1* suppress activated Gq. (A) The *mek-2(ku114)* mutation suppresses the exaggerated waveform of *egl-30*(*tg26*). N ≥ 14, *** *P* < 0.001, one-way ANOVA with Bonferroni’s *post hoc* test. (B) The *mpk-1(oz140)* mutation suppresses the exaggerated waveform of *egl-30*(*tg26*). N ≥ 12, *** *P* < 0.001, one-way ANOVA with Bonferroni’s *post hoc* test.

**Figure S2.**
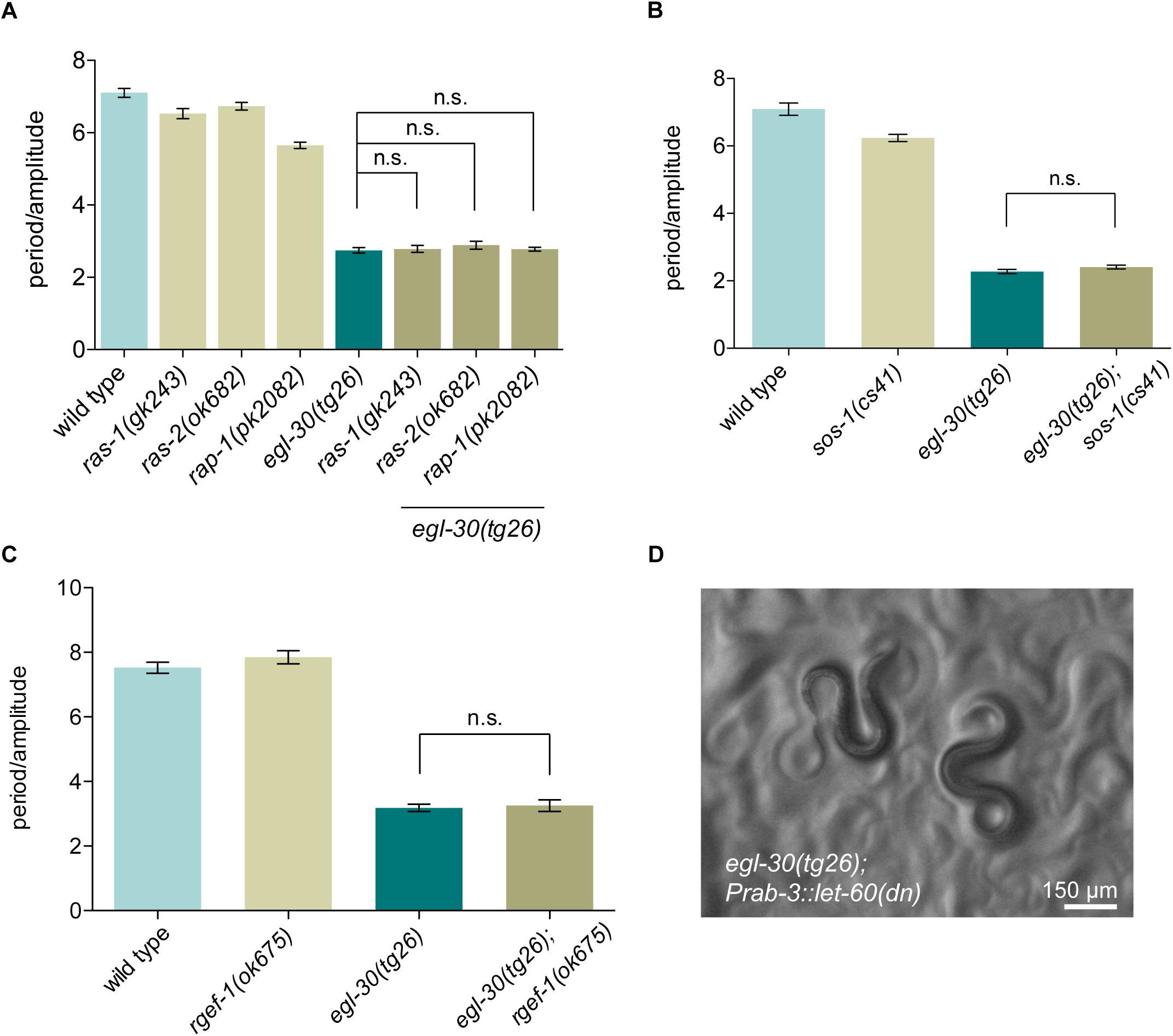
The exaggerated waveform of activated Gq does not depend on the RasGEFs *sos-1* and *rgef-1*, or the Ras family members *ras-1*, *ras-2*, and *rap-1*. (A) Deletion mutations *ras-1(gk243)* and *ras-2(ok682),* or the nonsense mutation *rap-1(pk2082)* do not suppress the waveform of the activated Gq mutant *egl-30(tg26)*. *N ≥* 18, n.s., not significant, one-way ANOVA with Bonferroni’s *post hoc* test. (B) Loss of the RasGEF *sos-1* does not suppress activated Gq waveform. Animals carrying the temperature sensitive *sos-1(cs41)* mutation were incubated for 24 hours at the non-permissive temperature of 25° and assayed for their waveform. *N ≥* 15, n.s., not significant, one-way ANOVA with Tukey’s *post hoc* test. (C) The RasGEF deletion mutation *rgef-1(ok675)* does not suppress the exaggerated waveform of activated Gq. *N =* 16, n.s., not significant, one-way ANOVA with Tukey’s *post hoc* test. (D) Neuronal expression of the dominant negative Ras mutant S17N (*yakEx158[Prab-3::let-60(S17N)]*) does not suppress the coiled posture of the activated Gq mutant *egl-30(tg26)*.

**Figure S3.**
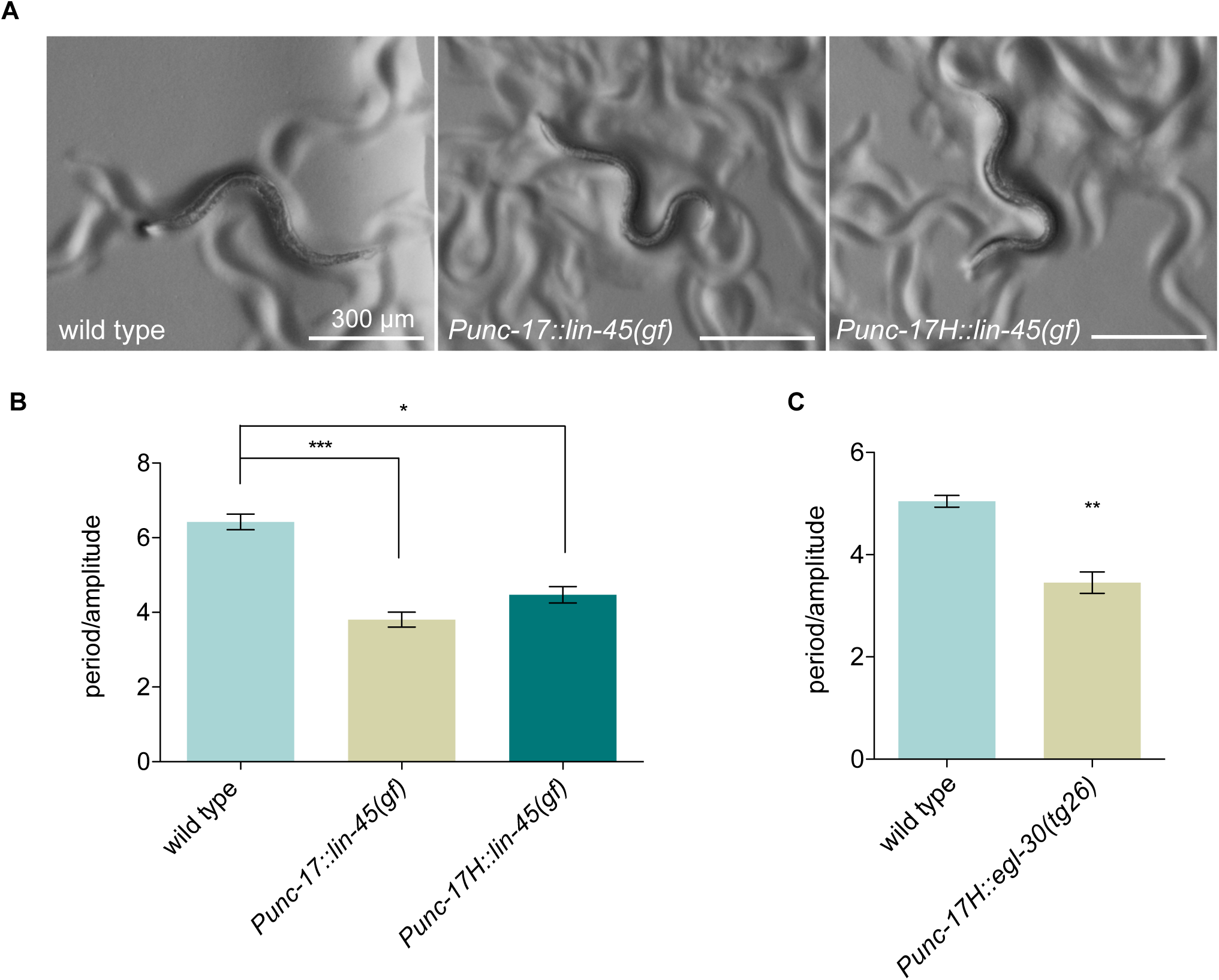
Expression of activated LIN-45/Raf or EGL-30/Gq in head acetylcholine neurons is sufficient to cause locomotion with an exaggerated waveform. (A, B) The *yakIs34[Punc-17::lin-45(gf)]* integrated array expressing activated *lin-45*(T626E/T629D) in all acetylcholine neurons and the *yakEx154[Punc-17H::lin-45(gf)]* extrachromosomal array expressed in head acetylcholine neurons both cause a coiled posture (A) and exaggerated waveform (B). *N ≥* 12, *** *P* < 0.001, * *P* < 0.05, Kruskal-Wallis test with Dunn’s *post hoc* test. (C) Expression of the activated Gq mutant *egl-30(tg26)* R243Q in head acetylcholine neurons (*Punc-17H::egl-30(tg26)*) causes an exaggerated waveform. N ≥ 7, ** P < 0.01, unpaired T test, two-tailed. Overexpression of activated Gq is toxic and these animals were slow growing and had reduced viability and fertility.

**Table S1.**
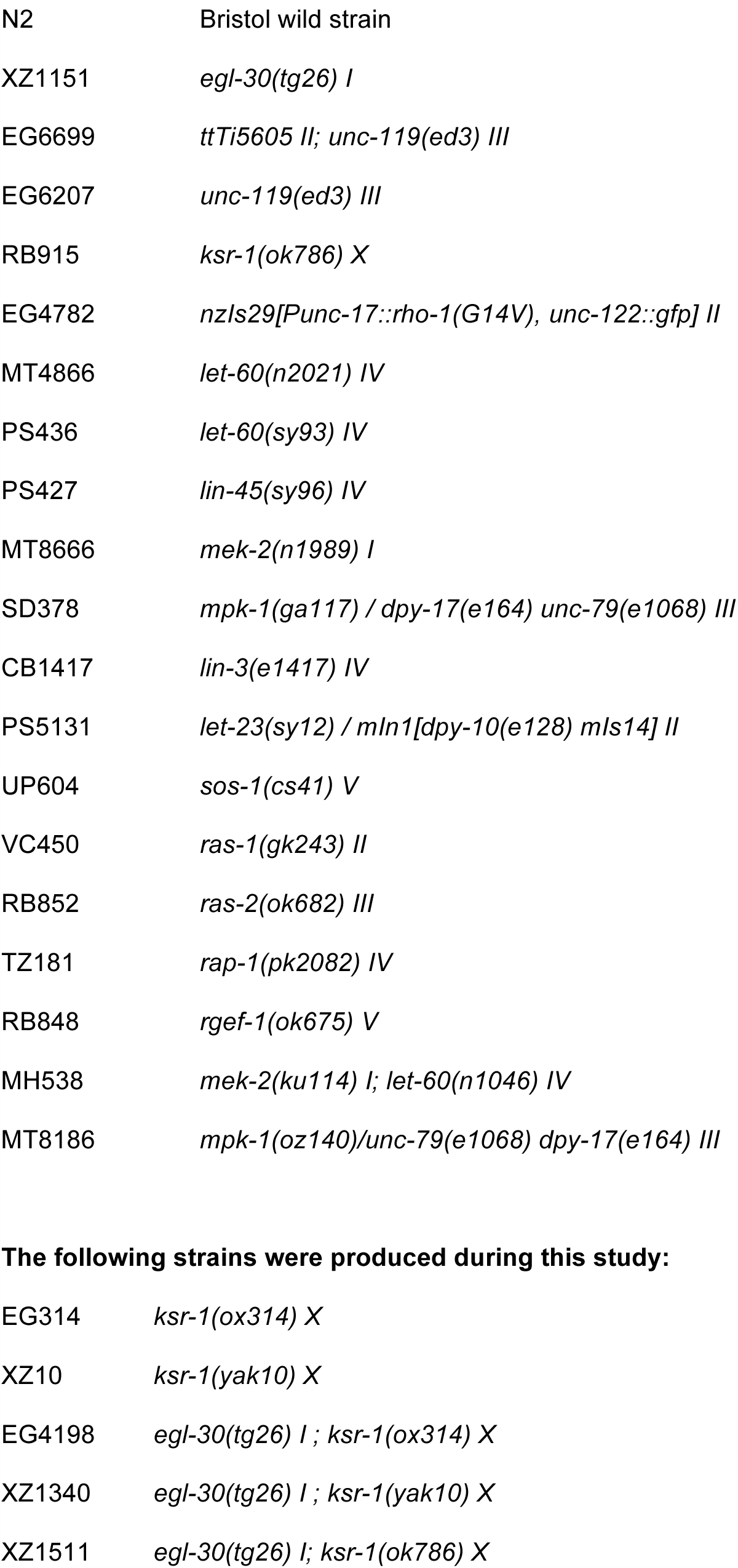
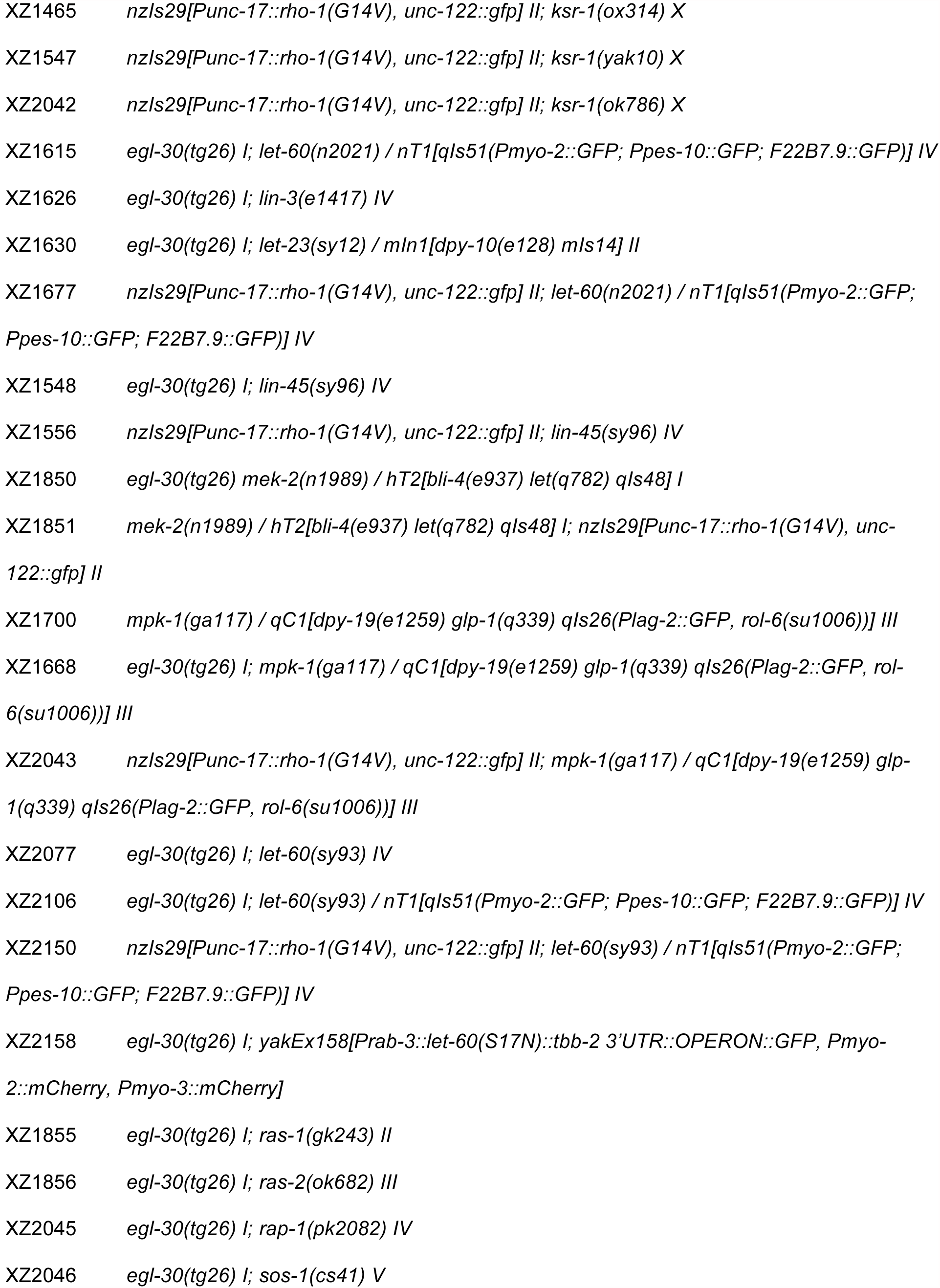
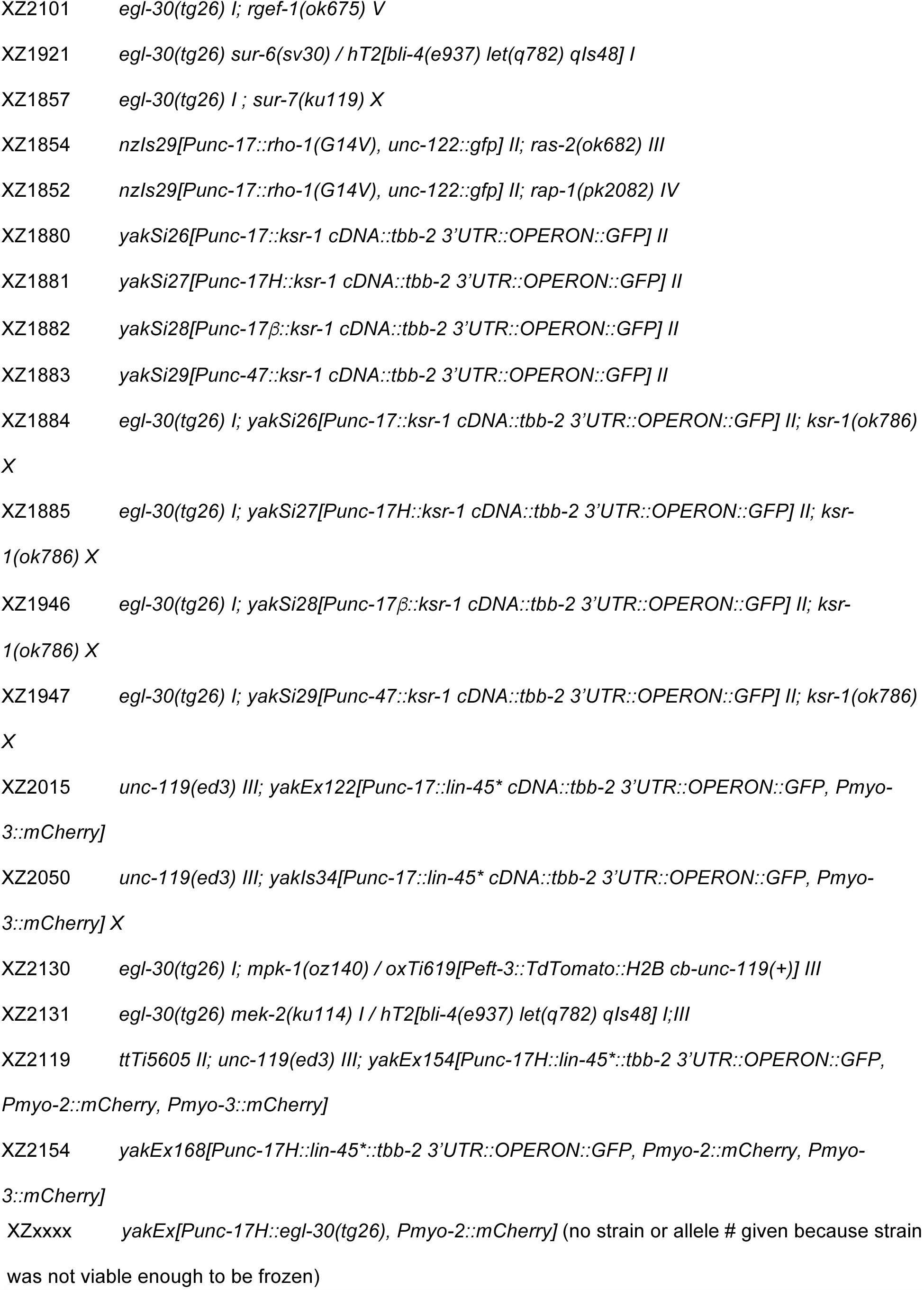
List of strains

**Table S2.**
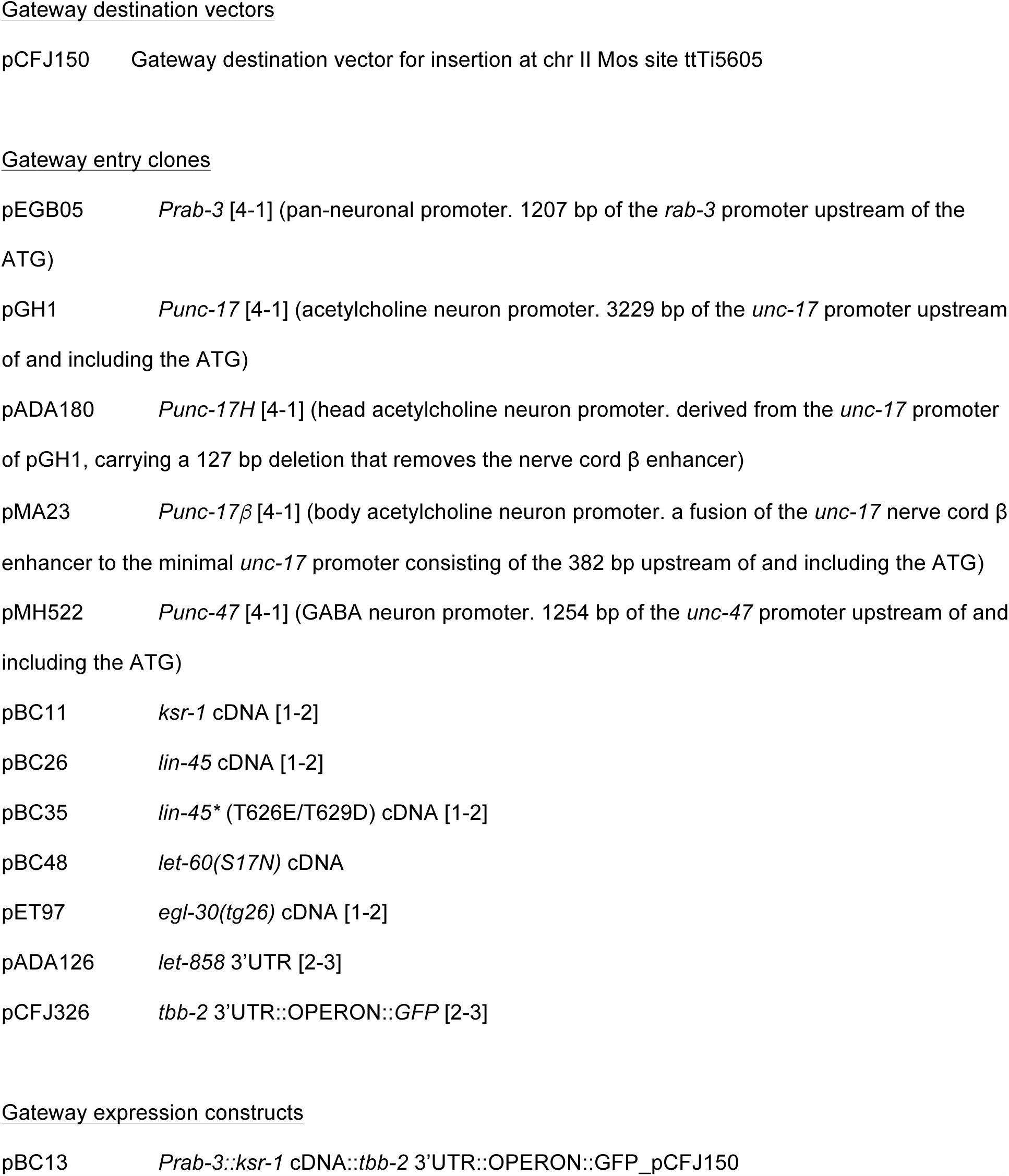
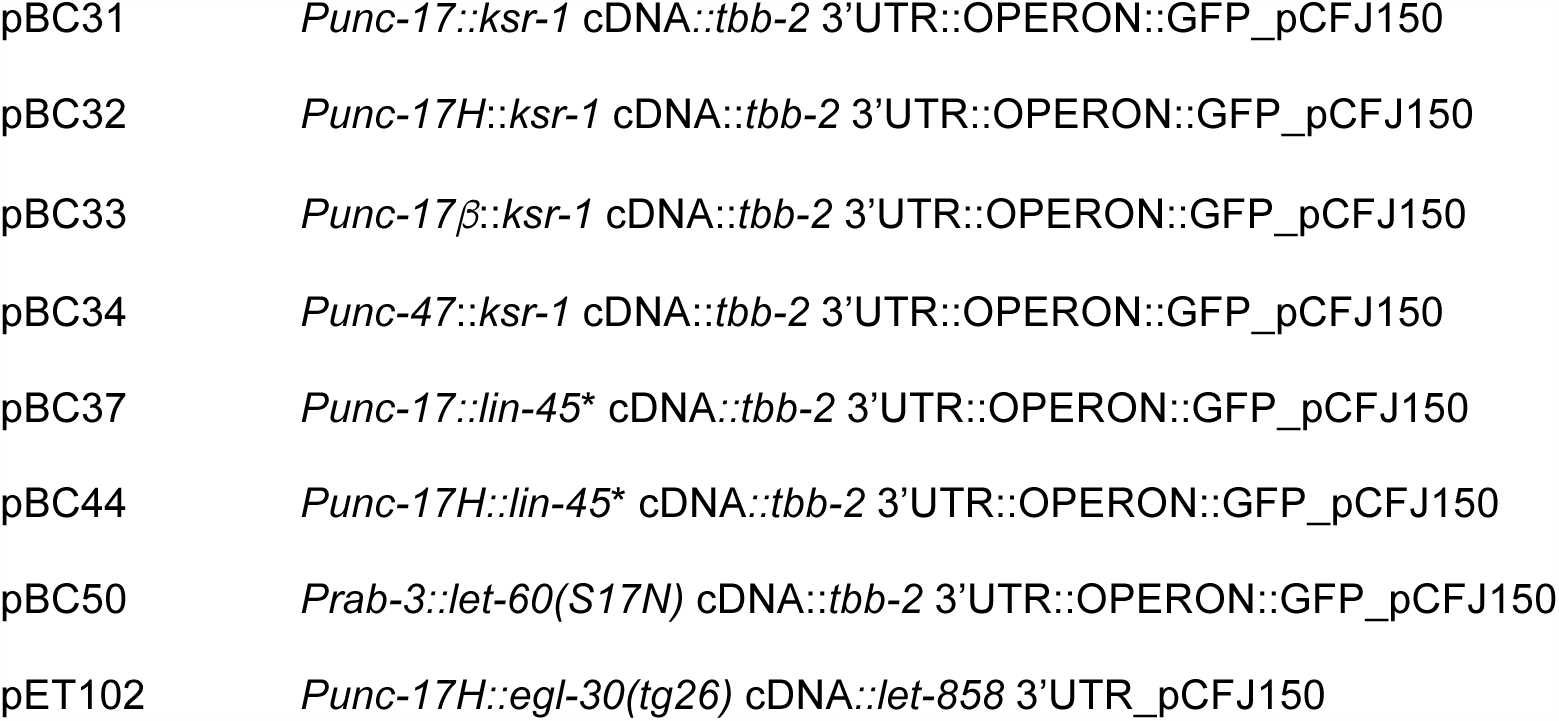
List of plasmids

**Table S3.**
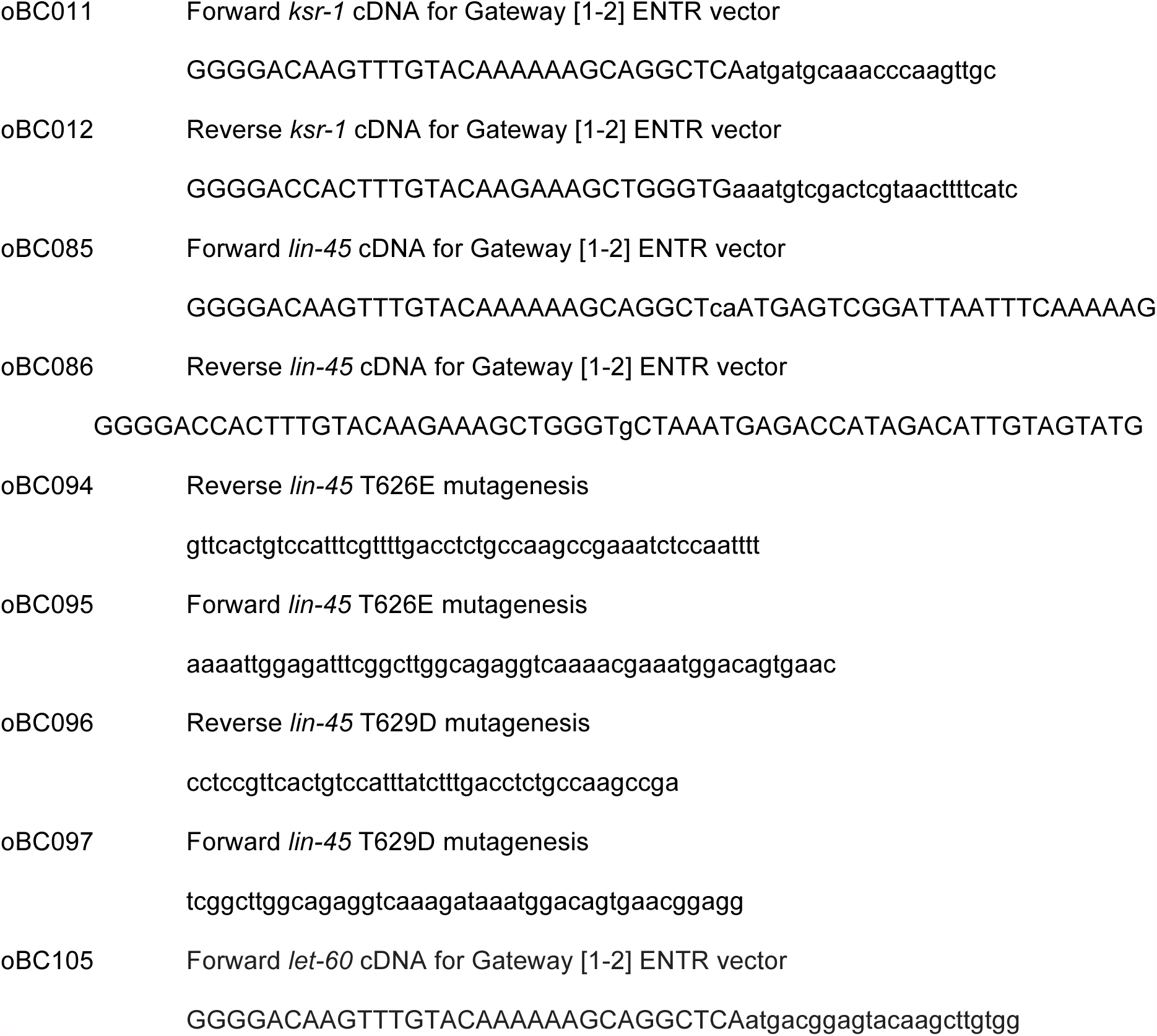
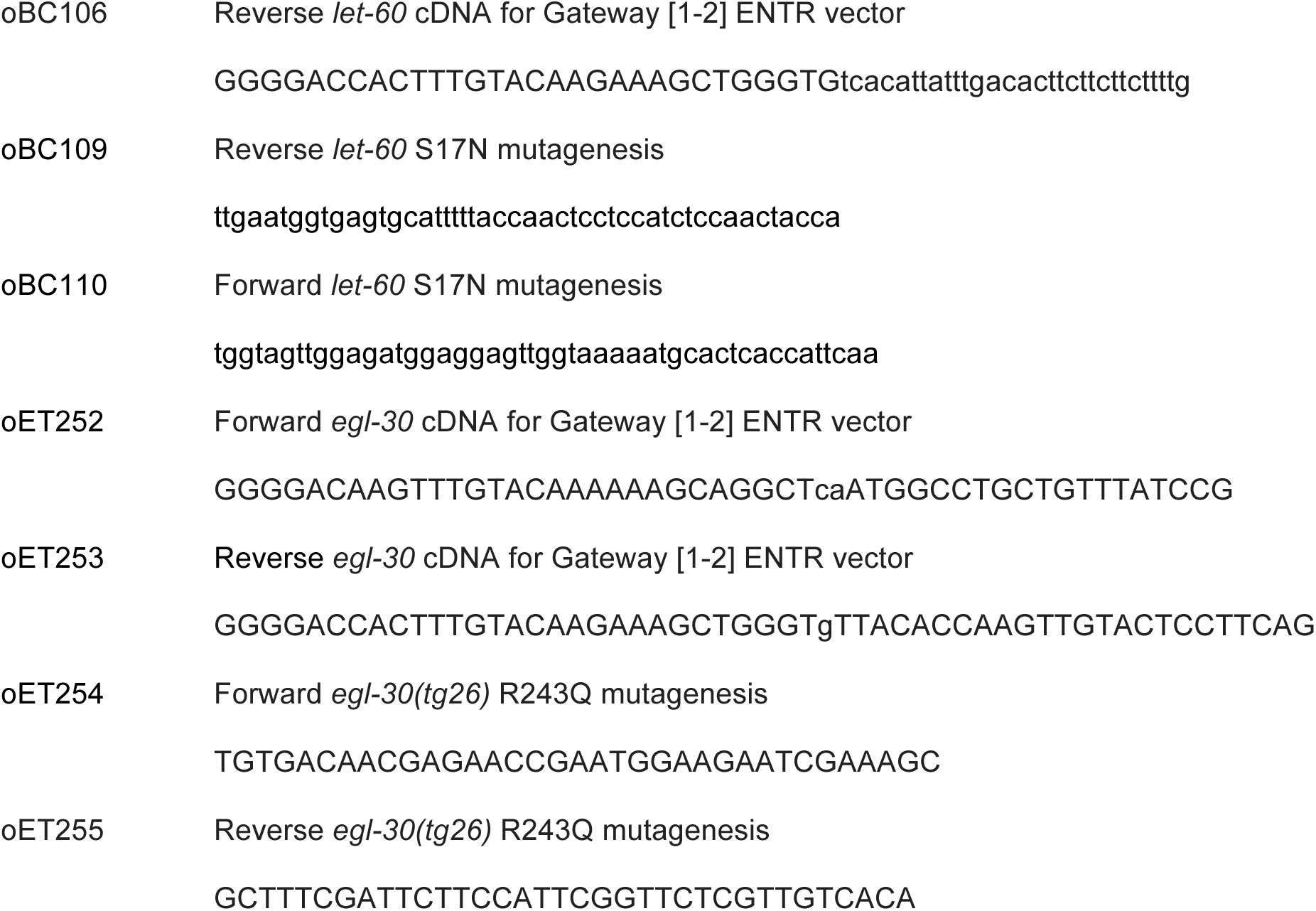
List of primers

